# Maturation and interhemispheric asymmetry in neurite density and orientation dispersion in early childhood

**DOI:** 10.1101/852764

**Authors:** Dennis Dimond, Stella Heo, Amanda Ip, Christiane S. Rohr, Ryann Tansey, Kirk Graff, Thijs Dhollander, Robert E. Smith, Catherine Lebel, Deborah Dewey, Alan Connelly, Signe Bray

## Abstract

**Background:** The brain’s white matter undergoes profound changes during early childhood, which are believed to underlie the rapid development of cognitive and behavioral skills during this period. Neurite density, and complexity of axonal projections, have been shown to change across the life span, though changes during early childhood are poorly characterized. Here, we utilize neurite orientation dispersion and density imaging (NODDI) to investigate maturational changes in tract-wise neurite density index (NDI) and orientation dispersion index (ODI) during early childhood. Additionally, we assess hemispheric asymmetry of tract-wise NDI and ODI values, and longitudinal changes.

**Methods:** Two sets of diffusion weighted images (DWI) with different diffusion-weighting were collected from 125 typically developing children scanned at baseline (N=125; age range=4.14-7.29; F/M=73/52), 6-month (N=8; F/M=8/0), and 12-month (N=52; F/M=39/13) timepoints. NODDI and template-based tractography using constrained spherical deconvolution were utilized to calculate NDI and ODI values for major white matter tracts. Mixed-effects models controlling for sex, handedness, and in-scanner head motion were utilized to assess developmental changes in tract-wise NDI and ODI. Paired t-tests were used to assess interhemispheric differences in tract-wise NDI and ODI values and longitudinal changes in cross-sectional and 12-month longitudinal analyses, respectively.

**Results:** Maturational increases in NDI were seen in all major white matter tracts, though we did not observe the expected tract-wise pattern of maturational rates (e.g. fast commissural/projection and slow frontal/temporal tract change). ODI did not change significantly with age in any tract. We observed higher cross-sectional NDI and ODI values in the right as compared to the left hemisphere for most tracts, but no hemispheric asymmetry for longitudinal changes.

**Conclusions:** These findings suggest that neurite density, but not orientation dispersion, increases with age during early childhood. In relation to NDI growth trends reported in infancy and late-childhood, our results suggest that early childhood may be a transitional period for neurite density maturation wherein commissural/projection fibers are approaching maturity, maturation in long range association fibers is increasing, and changes in limbic/frontal fibers remain modest. Rightward asymmetry in NDI and ODI values, but not longitudinal changes, suggests that rightward asymmetry of neurite density and orientation dispersion is established prior to age 4.

## 1 INTRODUCTION

Early childhood is an important maturational period characterized by rapid development of cognitive skills (Breckenridge et al., 2013; Burnett Heyes et al., 2012; Ferretti et al., 2008; Garon-Carrier et al., 2018; Montroy et al., 2016) and profound structural and functional changes in the brain (Reynolds et al., 2019a; Rohr et al., 2018; Sowell et al., 2004). Axonal fiber bundles (also referred to as “white matter tracts”) form the structural framework of the brain, facilitating rapid communication across brain networks and enabling distributed cognitive processing. Developmental changes in white matter properties are unsurprisingly linked to cognitive maturation (Bathelt et al., 2018; Jolles et al., 2016; Klarborg et al., 2013; Qiu et al., 2008; Van Eimeren et al., 2008), though these changes are understudied during early childhood, and trajectories for specific structural features are unclear.

Historically, neurodevelopmental studies have relied heavily on the diffusion tensor model (typically referred to as diffusion tensor imaging, or DTI) for insight into white matter structure. DTI studies suggest that white matter development is heterochronous, with changes in fractional anisotropy (FA) and mean diffusivity (MD) following a consistent tract-wise growth pattern; earlier maturation in commissural and projection fibers followed by association and then frontal-temporal fibers (Dubois et al., 2014; Lebel et al., 2012; Westlye et al., 2010). Developmental changes in DTI metrics are believed to reflect increases in axonal packing or myelination (Dubois et al., 2014; Lebel and Deoni, 2018; Mukherjee and McKinstry, 2006). However, each individual DTI metric is sensitive to a range of structural properties; they therefore lack the specificity needed to characterize changes in specific white matter features with age.

More recently, advanced biophysical diffusion models capable of estimating specific white matter properties (see Jelescu and Budde, 2017 for review) have been applied to study white matter development (Dean et al., 2017; Dimond et al., 2019a; Huber et al., 2019). One such model, called neurite orientation dispersion and density imaging (NODDI; Zhang et al., 2012), provides metrics sensitive to neurite/axonal density (neurite density index (NDI)) and the extent of orientational incoherence/dispersion amongst axonal projections within a voxel (orientation dispersion index (ODI)). Studies have shown that both NDI and ODI change across the lifespan and follow similar tract-wise patterns to changes in DTI metrics (Chang et al., 2015; Dean et al., 2017; Lebel and Beaulieu, 2011). Increases in NDI and ODI have been observed in early infancy (0-1 years old) (Batalle et al., 2017; Dean et al., 2017), though in childhood to adolescence (8-13 years old), increases are observed only for NDI (Mah et al., 2017); findings suggest ODI increases slow down sometime between infancy (1-3 years old) and childhood (Chang 2015; Jelescu 2015), only to speed back up in mid-adulthood (Chang 2015). In early childhood (ages 4-8 years old), we have previously found evidence of developmental increases in axonal density (Dimond et al., 2019a) quantified using 3-tissue constrained spherical deconvolution (Dhollander and Connelly, 2016) within the fixel-based analysis framework (FBA; Raffelt et al., 2017); these changes were consistent with established tract-wise growth patterns (Dubois et al., 2014; Lebel et al., 2012; Westlye et al., 2010), with more profound increases in projection and commissural fibers and no significant growth in frontal or temporal tracts. These observations suggest early childhood may be a dynamic maturational period in terms of neurite density and orientation dispersion, though NODDI has yet to be applied to investigate changes in early childhood specifically.

White matter maturation is known to occur asynchronously across the brain, though evidence of hemispheric asymmetries in white matter properties and developmental changes across the lifespan are largely inconsistent (Bonekamp et al., 2007; Carper et al., 2016; Dean et al., 2017; Takao et al., 2011; Thiebaut de Schotten et al., 2011). In infants and children, DTI metrics have been shown to have a leftward asymmetry, suggestive of faster left hemisphere maturation (Dubois et al., 2009; Krogsrud et al., 2016; Ratnarajah et al., 2013). In adolescents to adults, there is evidence of white matter differences across hemispheres (Bonekamp et al., 2007; Carper et al., 2016; Lebel and Beaulieu 2009; Takao et al., 2011; Thiebaut de Schotten et al., 2011), though most tracts show no asymmetry of age-related changes (Lebel and Beaulieu 2011; Takao et al., 2011). Taken together, this suggests that certain hemispheric differences in white matter structure may emerge early, and at some point between childhood and adulthood, rates of change become comparable across hemispheres. To our knowledge, only one study has explicitly tested for hemispheric differences in NDI and ODI values (Dean et al., 2017), finding evidence of both left and right regional asymmetries in both NDI and ODI in newborns. Whether neurite density and orientation dispersion values are asymmetric across hemispheres in early childhood, and if these properties demonstrate asymmetric developmental changes, has yet to be explored.

In this study, we utilized NODDI (Zhang et al., 2012) to investigate developmental changes in neurite density and orientation dispersion in early childhood, and to explore interhemispheric differences in cross-sectional NDI/ODI values and longitudinal NDI/ODI changes across major white matter tracts. Considering previous NODDI studies in infancy to adolescence (Dean et al., 2017; Mah et al., 2017), we hypothesized that NDI would increase with age in all major white matter tracts, while changes in ODI would be less widespread. In line with tract-wise developmental patterns seen for DTI metrics (Dubois et al., 2014; Lebel et al., 2012; Westlye et al., 2010), we further hypothesized that NDI and ODI changes would be most rapid in commissural and projection fibers, and slowest in frontal-temporal fibers. Finally, we hypothesized that NDI and ODI tract cross-sectional values and longitudinal changes would exhibit some degree of hemispheric asymmetry; though considering conflicting evidence for white matter hemispheric differences (Bonekamp et al., 2007; Takao et al., 2011; Thiebaut de Schotten et al., 2011), and scarce evidence for NDI and ODI (Dean et al., 2017), we did not make predictions about directionality.

## 2 MATERIALS AND METHODS

### 2.1 Participants

One-hundred and thirty-nine typically developing children (F/M=80/59) between the ages of 4.1 and 7.3 years were recruited from the Calgary area and underwent baseline MRI scanning. The study only included participants who were typically developing, with no history of neurological or psychiatric disorders and no contraindications to MR scanning. Fifty-five children returned for a 12-month follow-up (F/M=41/14), 10 of whom also participated in a 6-month follow-up (F/M=10/0). Twenty data sets were excluded from the study due to incomplete or poor data quality (described in more detail below). The final sample therefore consisted of 184 data sets collected from 125 participants at baseline (N=125; age range=4.14-7.29, mean=5.60, SD=0.81; F/M=73/52), 6-month follow-up (N=8; age range=4.61-7.01, mean=5.61, SD=0.78; F/M=8/0), and 12-month follow-up (N=52; age range=5.13-7.89, mean=6.63, SD=0.76; F/M=39/13). The study was approved by the Conjoint Health and Research Ethics Board at the University of Calgary. Informed consent was obtained from the parents and informed assent from the participants.

### 2.2 Data acquisition and motion prevention

Data were collected at the Alberta Children’s Hospital and included diffusion magnetic resonance imaging data, as well as parent-reported handedness. MRI data were acquired on a 3T GE MR750w (Waukesha, WI) scanner using a 32-channel head coil. Two separate sets of diffusion-weighted images (DWIs) were acquired at b=1000s/mm^2^ and b=2000s/mm^2^ respectively using 2D spin-echo EPI sequences (45 diffusion-weighted directions, 3 non-DWI (*b*=0) images, 2.5mm^3^ isotropic voxels, 45 slices, 23×23cm FOV, TE=86.2ms, TR=10s).

Strategies to prevent in-scanner head motion were implemented leading up to and during MRI scanning at each session. In the week leading up to MRI scanning, participants were introduced to the scanning environment via a mock-scan and parents were asked to have their child practice lying still at home. During MRI scanning, padding was inserted between each side of the participant’s head and the head-coil. Participants were permitted to watch a movie of their choosing during scanning, and we provided audible reminders to the participants to try to “remain perfectly still” if we observed the participant moving in the scanner.

### 2.3 Preprocessing and motion assessment

To mitigate the effects of in-scanner head motion, DWI preprocessing made use of recent advances in motion-correction (Andersson and Sotiropoulos, 2016), included replacement of slice-wise signal dropout (Andersson et al., 2016) and within-volume motion correction (Andersson et al., 2017) implemented in FSL (Jenkinson et al., 2012). DWI preprocessing was performed separately for the b=1000s/mm^2^ and b=2000s/mm^2^ shells. DWI volumes for which more than 20% of slices were classified as containing signal dropout were removed from the data, and scans for which more than 10% of volumes were removed were excluded from the study entirely. Since at least two diffusion weighted shells are required for the NODDI model, datasets were excluded if either the b=1000s/mm^2^ or b=2000s/mm^2^ shell were excluded. Following this approach, 15 datasets were excluded from the study. To quantify and control for the potential influence of head motion on outcome measures, we calculated the total number of signal dropout slices per dataset for use as a covariate in statistical analyses. Within-volume and between-volume motion parameters were also calculated. The total number of signal dropout slices was selected as our main motion covariate because it captures both the extent of motion in the data as well as the extent of signal dropout replacement, and had one of the strongest associations with age. Motion characteristics for this data set and associations with age are provided in Supplementary Table 1; none of the motion metrics were significantly correlated with age. As a final step, we visually inspected the preprocessed data to ensure within-volume motion artifacts had been sufficiently removed, and that the whole cerebrum was within the field of view. In 5 scans, substantial in-scanner head movement had resulted in incomplete temporal lobe coverage in several diffusion volumes. These 5 datasets were therefore excluded from the study.

### 2.4 White Matter template generation

Major white matter tracts were delineated from a study-specific fiber orientation distribution (FOD) template created in *MRtrix3* (Tournier et al., 2019). To do this, b=2000s/mm^2^ images were first upsampled to 1.25mm^3^ isotropic voxel size, which can potentially benefit template creation and template-based tractography (Dyrby et al., 2014). An FOD image was then generated for each data set using single-shell 3-tissue constrained spherical deconvolution (SS3T-CSD) (Dhollander and Connelly, 2016), using a group averaged response function for each tissue type (white matter, grey matter and cerebrospinal fluid) (Dhollander et al., 2016; Raffelt et al., 2012); SS3T-CSD was carried out using *MRtrix3*Tissue (https://3tissue.github.io/), a fork of *MRtrix3* (Tournier et al., 2019) (for interested readers, this method is described in full detail in Dhollander and Connelly, 2016, and is also summarized in Dimond et al., 2019a). FOD images from 40 (F/M=20/20) participants – selected such that the full study sample age range was represented with preferential selection of images with fewer number of total dropout slices – were utilized to generate the study-specific FOD template via an iterative registration and normalization approach (Raffelt et al., 2011).

### 2.5 Template-based tractography

Utilizing the study-specific FOD template, we defined a single set of tracts for use in calculation of mean tract NDI and ODI values for all participant scans. Probabilistic tractography (step size=0.625mm, max angle between steps=22.5°, min/max fiber length=10mm/250mm, cut-off FOD amplitude=0.1) and random seeding throughout the FOD template were performed to create a whole-brain tractogram with 20 million streamlines. Global reconstruction bias reduction using spherical-deconvolution informed filtering of tractograms (SIFT) (Smith et al., 2013) was then applied to reduce the tractogram to 2 million streamlines. From this reduced tractogram, we extracted the following major white matter tracts of interest, which we divide approximately into 3 groups for visualization purposes: 1) the corticospinal tracts, superior longitudinal, inferior frontal-occipital, and inferior longitudinal fasciculi (Figure 1); 2) the arcuate fasciculi, cingulum bundles, fornix bundles, and uncinate fasciculi (Figure 2); 3) the genu, body and splenium of the corpus callosum and their projections to the cortex (Figure 3). Tracts were extracted using manually-defined inclusion and exclusion ROIs with placement consistent with those used in Dimond et al., 2019. For each tract we generated a binary voxel mask that included only voxels through which at least 5 streamlines passed. To assess global brain changes, we also generated a global white matter voxel mask by applying the same streamline threshold to the 2 million streamline tractogram.

**Figure 1.**
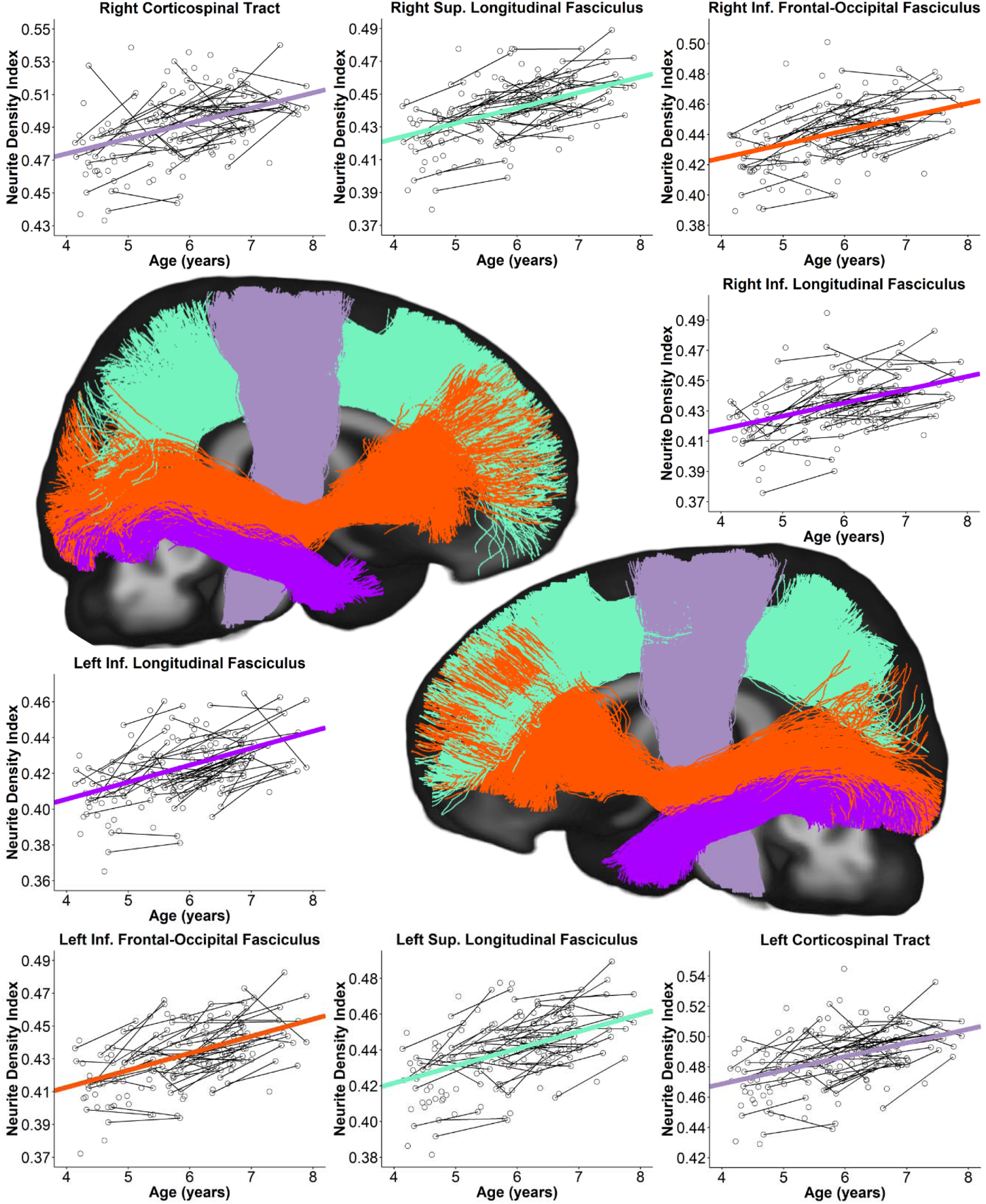
Scatterplots of white matter tract NDI values from ages 4 to 8 years. Scatterplots of individual NDI values (open circles) for the superior longitudinal, inferior frontal-occipital, and inferior longitudinal fasciculi and corticospinal tracts overlaid with trendline trajectories (bold coloured lines). Scans collected from the same participant are indicated by a black line connecting the open circles. White matter tracts are visualized as 3D streamlines overlaid onto a 2D mid-sagittal slice of the study-specific FOD template, and colour coordinated to match the scatterplot trendline trajectories. Rates of change (i.e. trendline slopes) for all tracts are provided in Table 1.

**Figure 2.**
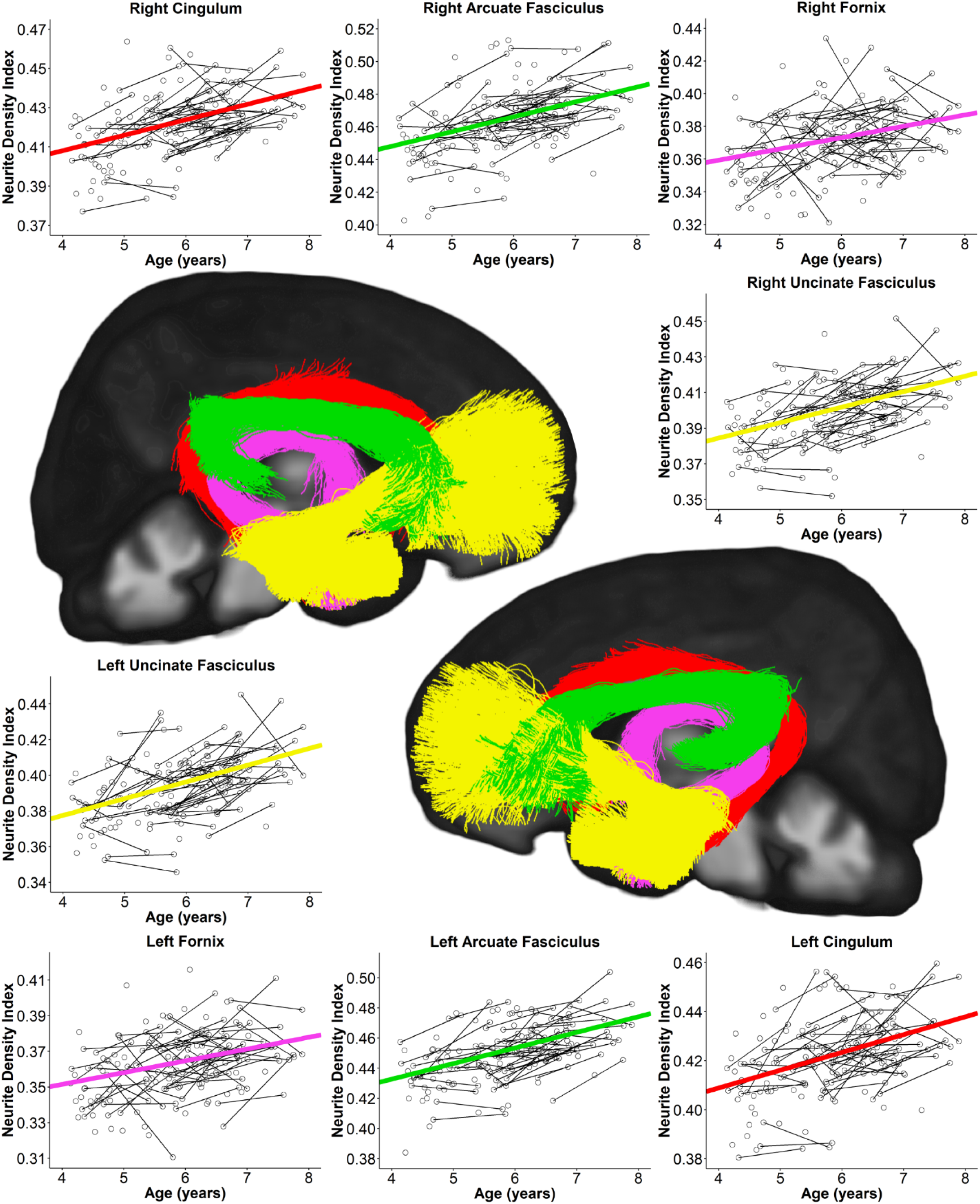
Scatterplots of white matter tract NDI values from ages 4 to 8 years. Scatterplots of individual NDI values (open circles) for the cingulum and fornix bundles, and uncinate and arcuate fasciculi overlaid with trendline trajectories (bold coloured lines). Scans collected from the same participant are indicated by a black line connecting the open circles. White matter tracts are visualized as 3D streamlines overlaid onto a 2D mid-sagittal slice of the study-specific FOD template, and colour coordinated to match the scatterplot trendline trajectories. Rates of change (i.e. trendline slopes) for all tracts are provided in Table 1.

**Figure 3.**
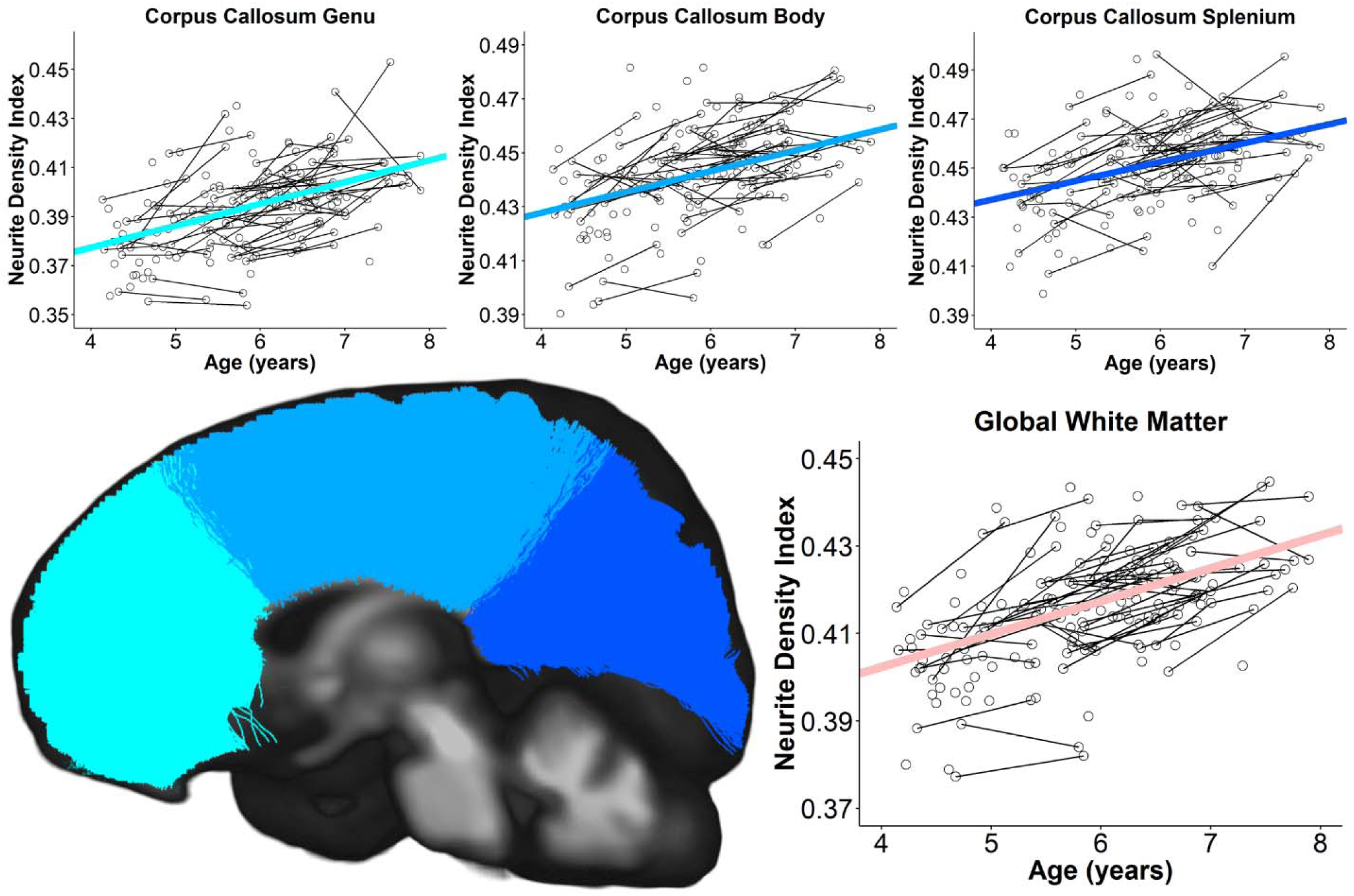
Scatterplots of commissural and global white matter NDI values from ages 4 to 8 years. Scatterplots of individual NDI values (open circles) for the corpus callosum segments and global white matter mask overlaid with trendline trajectories (bold coloured lines). Scans collected from the same participant are indicated by a black line connecting the open circles. White matter tracts are visualized as 3D streamlines overlaid onto a 2D mid-sagittal slice of the study-specific FOD template, and colour coordinated to match the scatterplot trendline trajectories. As the global white matter mask encompasses all voxels with at least some white matter partial volume, no global white matter mask streamlines are shown. Rates of change (i.e. trendline slopes) for all tracts are provided in Table 1.

**Table 1:** Developmental trajectory values of tract-wise NDI. NDI increased significantly from ages 4 to 8 in all tracts.

| | Rate of Change<br>($\Delta$/year) | Predicted Value<br>at Age 4<br>(NDI$\times 10^{-1}$) | Predicted Value<br>at Age 8<br>(NDI$\times 10^{-1}$) | Percent Increase<br>from Ages 4-8 (%) |
| --- | --- | --- | --- | --- |
| Right Corticospinal Tract | $9.3 \times 10^{-3}$ | 4.7 | 5.1 | 7.8 |
| Left Corticospinal Tract | $9.1 \times 10^{-3}$ | 4.7 | 5.1 | 7.7 |
| Right Sup. Longitudinal Fasciculus | $9.4 \times 10^{-3}$ | 4.2 | 4.6 | 8.9 |
| Left Sup. Longitudinal Fasciculus | $9.6 \times 10^{-3}$ | 4.2 | 4.6 | 9.1 |
| Right Inf. Frontal-Occipital Fasciculus | $9.0 \times 10^{-3}$ | 4.2 | 4.6 | 8.5 |
| Left Inf. Frontal-Occipital Fasciculus | $1.0 \times 10^{-2}$ | 4.1 | 4.5 | 10.1 |
| Right Inf. Longitudinal Fasciculus | $8.8 \times 10^{-3}$ | 4.2 | 4.5 | 8.4 |
| Left Inf. Longitudinal Fasciculus | $9.5 \times 10^{-3}$ | 4.1 | 4.4 | 9.4 |
| Right Arcuate Fasciculus | $9.1 \times 10^{-3}$ | 4.5 | 4.8 | 8.1 |
| Left Arcuate Fasciculus | $1.0 \times 10^{-2}$ | 4.3 | 4.7 | 9.6 |
| Right Cingulum | $7.9 \times 10^{-3}$ | 4.1 | 4.4 | 7.7 |
| Left Cingulum | $7.3 \times 10^{-3}$ | 4.1 | 4.4 | 7.1 |
| Right Fornix | $6.9 \times 10^{-3}$ | 3.6 | 3.9 | 7.7 |
| Left Fornix | $6.6 \times 10^{-3}$ | 3.5 | 3.8 | 7.5 |
| Right Uncinate Fasciculus | $8.7 \times 10^{-3}$ | 3.8 | 4.2 | 9.0 |
| Left Uncinate Fasciculus | $9.4 \times 10^{-3}$ | 3.8 | 4.2 | 10.0 |
| Corpus Callosum Genu | $8.9 \times 10^{-3}$ | 3.8 | 4.1 | 9.4 |
| Corpus Callosum Body | $7.7 \times 10^{-3}$ | 4.3 | 4.6 | 7.2 |
| Corpus Callosum Splenium | $7.7 \times 10^{-3}$ | 4.4 | 4.7 | 7.1 |
| Global White Matter Mask | $7.5 \times 10^{-3}$ | 4.0 | 4.3 | 7.5 |

### 2.6 NODDI processing

The NODDI toolbox (https://www.nitrc.org/projects/noddi_toolbox) in MATLAB 2016b (Mathworks Inc., Natick, MA, USA) was utilized to calculate NDI and ODI maps for each data set after rigid-body registration and combining the b=1000s/mm^2^ shell with the b=2000s/mm^2^ shell. The NDI and ODI images were then warped to FOD template space by applying the deformation field generated by registering each participant’s FODs to the FOD template using *MRtrix3*. For the voxel masks in template space corresponding to each tract of interest, we calculated the mean values of both NDI and ODI for each participant.

Both NDI and ODI values range from 0-1. NDI reflects the fraction of the diffusion signal that is attributed to the intra-neurite compartment, as opposed to the extra-neurite compartment: a large NDI value (approaching 1) indicates that almost all signal comes from the intra-neurite compartment, suggestive of large and/or densely packed neurites or axons; a small NDI value (approaching 0) indicates that almost no signal comes from the intra-neurite compartment, suggestive of small and/or sparse neurites or axons. ODI reflects the extent of dispersion in estimated neurite orientations: a large ODI value (approaching 1) indicates near-isotropic distribution of neurite orientations; a small value (approaching 0) indicates that all orientations within a voxel are nearly parallel (Zhang et al., 2012). Both NDI and ODI are dimensionless measures, and therefore have no units.

### 2.7 Statistics: age associations

Linear mixed effects models performed using *lme4* (Bates et al., 2017) in R (R Core Team, 2014) were utilized to assess changes in tract-wise NDI and ODI with age. A linear mixed effects model is a regression model that can account for unequal numbers of data points across participants with unequal time intervals between data points, making it an ideal model for assessing neurodevelopmental growth in cross-lagged longitudinal studies (Genc et al., 2019; Lebel and Beaulieu, 2011; Westerhausen et al., 2016). Here we utilized mixed effects models to account for repeated measures within individual participants, and unequal time intervals between scans, by modelling each participant’s intercept as a random effect. Models were run separately for NDI and ODI, with age, handedness, sex, and total number of dropout slices as fixed effects, and participant-specific intercepts as a random effects. The total number of dropout slices was not significantly correlated with age in the full sample (r=-0.129, uncorrected p-value=0.081), though was included as a nuisance covariate to account for potential influence on outcome measures. P-values for t-statistics were obtained using the Satterthwaite’s degrees of freedom method (Hrong-Tai Fai and Cornelius, 2007), as implemented in *lmerTest* (Kuznetsova et al., 2017). False discovery rate (FDR) was utilized to control for multiple comparisons within each NODDI metric (corrected for 20 regions: 19 tracts + the global white matter mask), with statistical inferences drawn at *p*<0.05. Models were assessed both with and without outliers (defined as being greater than the 3^rd^ quartile + 1.5*(interquartile range), or less than the first quartile −1.5*(interquartile range)), though exclusion of outliers did not affect inferences drawn from the results, and so the results shown in this study are inclusive of all data. Here we report the rates of change (slope of the age regression, i.e. the beta value), predicted intercept values at ages 4 and 8, and the tract-wise percentage increases from ages 4-8 (calculated by dividing the difference between the predicted values at ages 8 and 4 years by the predicted value at age 4).

### 2.8 Statistics: interhemispheric asymmetries

In addition to tract-wise maturational rates, we assessed interhemispheric differences in NDI and ODI values as well as interhemispheric differences in longitudinal changes for lateralized tracts (i.e. all segmented tracts except for the corpus callosum segments and the global white matter mask). Differences in left vs. right tract values were assessed via paired t-tests, conducted cross-sectionally in a sample generated by selecting the scan with the least motion for each participant (N=125; age range=4.21-7.76, mean=5.79, SD=0.89; F/M=73/52). Interhemispheric differences in longitudinal change were assessed by subtracting baseline tract values from 12-month follow-up values in participants who had data from two timepoints (N=52; baseline: age range=4.14-6.88, mean=5.57, SD=0.77; F/M=39/13; 12-month follow-up: age range=5.13-7.89, mean=6.63, SD=0.76; F/M=39/13), and then conducting paired t-tests to compare changes in left vs. right hemisphere tracts. Analyses were conducted both with and without outliers (as defined above), though inferences drawn from the results were unchanged with exclusion of outliers, and only analyses including outliers are reported here. FDR with statistical inference set at *p*<0.05 was utilized to correct for multiple comparisons within cross-sectional and longitudinal analyses for each metric separately, controlling for the 16 lateralized tracts per analysis. For these analyses we present: mean values/longitudinal changes in left and right hemisphere tracts; differences in mean values between hemispheres; and interhemispheric percentage differences, calculated as the difference between left and right tract values or longitudinal changes thereof divided by the mean of left and right hemisphere values ((right hemisphere – left hemisphere) / mean(left hemisphere, right hemisphere)).

## 3 RESULTS

### 3.1 Rates of change and estimated intercepts

Significant increases in NDI were found in all white matter tracts, with rates of change of 6.6×10^-3^ to 1.0×10^-2^/year, and percent NDI increases from 7.1-10.1% from ages 4 to 8 (Table 1). Tract-wise scatter plots overlaid with estimated trend lines suggest linear fits were appropriate within the study age range (Figures 1-3). Faster increases were observed in the left inferior frontal-occipital fasciculus and left arcuate fasciculus, while slower increases were observed in the left and right fornix bundles (Figure 4). Larger percentage increases were seen in the left inferior frontal-occipital fasciculus and left uncinate fasciculus, while smaller percentage increases were seen in the splenium and left cingulum. NDI of the global white matter mask increased at a rate of 7.5×10^-3^/year (Figure 3), with a total increase of 7.5% from ages 4 to 8 (Table 1). Estimated age 4 intercept values for NDI (Table 1) were largest in the bilateral corticospinal tracts, bilateral arcuate fasciculi, and splenium and body of the corpus callosum, and smallest in the bilateral fornix bundles, genu of the corpus callosum, and bilateral uncinate fasciculi. Tract-wise predicted age 8 intercepts followed the same pattern. As seen in Figure 4, we did not observe the expected tract-wise pattern of rates of change in NDI across tracts (i.e. faster commissural/projection maturation and slower frontal/temporal growth (Dubois et al., 2014; Lebel et al., 2012; Westlye et al., 2010)).

**Figure 4.**
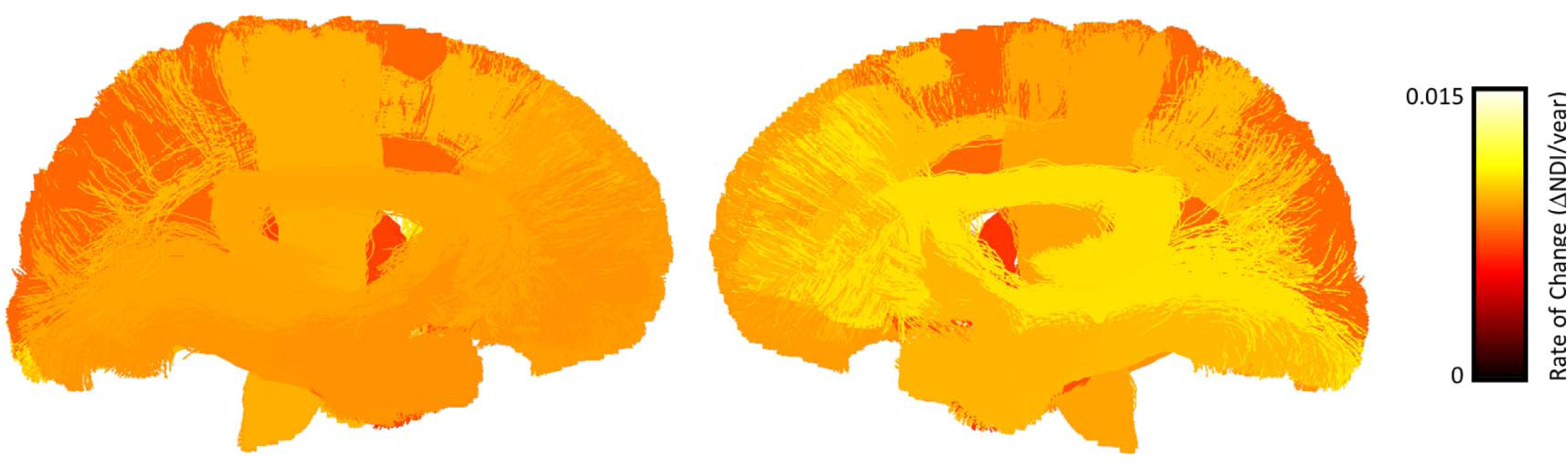
Anatomical visualization of tract-wise rates of change in NDI. Tract-wise rates of change in NDI are shown for all tracts assessed in the study, excluding the global white matter mask. Streamlines corresponding to each tract are coloured according to the rate of change of NDI observed in that tract.

ODI did not increase or decrease significantly in any of the assessed white matter tracts (Supplementary Table 2). Estimated age 4 intercept values for ODI were largest in the bilateral inferior longitudinal fasciculi, global white matter mask, and right uncinate fasciculus, and smallest in the bilateral fornix bundles, corticospinal tracts, and inferior frontal-occipital fasciculi. Estimated intercepts at age 8 demonstrated generally the same pattern, reflecting no significant change in relative ODI values across tracts.

### 3.2 Interhemispheric differences

Small but significant cross-sectional interhemispheric differences in NDI were found in all tracts except for the cingulum, with lower NDI in left as compared to right hemisphere tracts (Table 2). Amongst tracts with significant interhemispheric NDI differences, left hemisphere tract values were on average 8.2×10^-3^, or 1.9% lower than right hemisphere tract values. For ODI, cross-sectional tract values were significantly lower in the left as compared to the right hemisphere for all tracts. Similar to NDI, interhemispheric differences in ODI were small, with an average difference of 6.0×10^-3^, or 1.9% across tracts (Table 3). We found no significant interhemispheric differences in longitudinal changes for NDI or ODI.

**Table 2:** Cross-sectional interhemispheric differences in NDI. Left and right hemisphere tract values, and absolute and relative interhemispheric differences in cross-sectional NDI. * indicates tracts for which there was a significant difference in cross-sectional NDI values across hemispheres.

| | <u>Absolute Values</u><br>(NDI$\times 10^{-1}$) | | <u>Absolute Difference</u> | <u>Relative Difference</u> | <u>Effect Size</u> |
| --- | --- | --- | --- | --- | --- |
|  | **Right** | **Left** | **Right – Left** | **Percent (%)** | **Cohen's *d*** |
| Corticospinal Tract | 4.9 | 4.8 | $5.6 \times 10^{-3}$ | 1.1 | 0.8* |
| Sup. Longitudinal Fasciculus | 4.4 | 4.4 | $1.1 \times 10^{-3}$ | 0.2 | 0.2* |
| Inf. Frontal-Occipital Fasciculus | 4.4 | 4.3 | $9.9 \times 10^{-3}$ | 2.3 | 1.1* |
| Inf. Longitudinal Fasciculus | 4.3 | 4.2 | $1.1 \times 10^{-2}$ | 2.7 | 1.2* |
| Arcuate Fasciculus | 4.6 | 4.5 | $1.5 \times 10^{-2}$ | 3.2 | 1.6* |
| Cingulum | 4.2 | 4.2 | $2.5 \times 10^{-4}$ | 0.1 | 0.0 |
| Fornix | 3.7 | 3.6 | $8.7 \times 10^{-3}$ | 2.4 | 0.9* |
| Uncinate Fasciculus | 4.0 | 3.9 | $6.3 \times 10^{-3}$ | 1.6 | 1.0* |

**Table 3:** Cross-sectional interhemispheric differences in ODI. Left and right hemisphere tract values, and absolute and relative interhemispheric differences in cross-sectional ODI. Hemispheric ODI cross-sectional values were significantly different for all tracts.

| | <u>Absolute Values</u><br>(ODI$\times 10^{-1}$) | | <u>Absolute Difference</u> | <u>Relative Difference</u> | <u>Effect Size</u> |
| --- | --- | --- | --- | --- | --- |
|  | **Right** | **Left** | **Right – Left** | **Percent (%)** | **Cohen's *d*** |
| Corticospinal Tract | 3.0 | 2.9 | $5.6 \times 10^{-3}$ | 1.9 | 0.9 |
| Sup. Longitudinal Fasciculus | 3.2 | 3.2 | $5.5 \times 10^{-3}$ | 1.7 | 0.8 |
| Inf. Frontal-Occipital Fasciculus | 3.1 | 3.1 | $1.7 \times 10^{-3}$ | 0.5 | 0.2 |
| Inf. Longitudinal Fasciculus | 4.1 | 4.0 | $3.3 \times 10^{-3}$ | 0.8 | 0.2 |
| Arcuate Fasciculus | 3.5 | 3.5 | $3.2 \times 10^{-3}$ | 0.9 | 0.3 |
| Cingulum | 3.5 | 3.4 | $1.2 \times 10^{-2}$ | 3.5 | 1.0 |
| Fornix | 2.4 | 2.3 | $6.4 \times 10^{-3}$ | 2.8 | 0.7 |
| Uncinate Fasciculus | 3.6 | 3.5 | $1.0 \times 10^{-2}$ | 2.9 | 1.1 |

## 4 DISCUSSION

The aim of the current study was to characterize developmental trends in tract-wise neurite density and orientation dispersion in early childhood and determine if cross-sectional values and/or longitudinal changes of such differ between left and right hemispheres. We found developmental increases in NDI, but no change in ODI, in all major white matter tracts, as well as in the global white matter mask. Contrary to our hypothesis, NDI increases did not follow expected tract-wise growth trends (i.e. faster commissural/projection fiber maturation and slower frontal/temporal tract maturation). Instead, tracts matured at relatively similar rates across the brain with no clear tract-wise pattern of maturation. Interestingly, we found a rightward asymmetry (right>left) in cross-sectional NDI and ODI tract values for almost all tracts, though no significant interhemispheric differences in longitudinal growth were observed. These findings add to our understanding of white matter development during early childhood, and provide insight into tract-wise trends in neurite density and orientation incoherence with age. Below we discuss the implications of our findings in relation to the NODDI and DTI literature, evidence of axonal/neurite density maturation from other biophysical diffusion models and histology, and the functional relevance of these changes.

### 4.1 Development of neurite density and orientation dispersion

Utilizing NODDI (Zhang et al., 2012), several studies have demonstrated substantial white matter microstructural changes across the lifespan, characterized by approximately logarithmic increases in NDI from birth to adulthood (Batalle et al., 2017; Chang et al., 2015; Dean et al., 2017; Genc et al., 2017a; Huber et al., 2019; Jelescu et al., 2015; Mah et al., 2017; Young et al., 2019), and delayed exponential increases in ODI (Chang et al., 2015), with marginal increases in ODI during early infancy (0-1 years) (Batalle et al., 2017; Dean et al., 2017), no change during childhood and adolescence (Mah et al., 2017; Young et al., 2019), and more rapid increases in mid-late adulthood (Billiet et al., 2015; Chang et al., 2015; Kodiweera et al., 2016; Ota et al., 2017).

Our findings build on this literature by bridging the gap between infant-toddler studies and late childhood. Consistent with trends in the literature, we find rapid NDI increases in all major white matter tracts, and no significant change in ODI during early childhood. In terms of estimated intercepts, at both ages 4 and 8, we find highest NDI values in the corticospinal tracts, splenium and body of the corpus callosum, and lowest values in the fornix bundles, uncinate fasciculi, and genu (inclusive of its frontal projections). ODI values at these ages demonstrate the inverse relationship, with highest values in frontal-temporal tracts (e.g. the uncinate and inferior longitudinal fasciculi) and lowest values in the corticospinal tracts. These findings are consistent with observations from newborns to adults (Chang et al., 2015; Dean et al., 2017; Kunz et al., 2014; Mah et al., 2017), suggesting that the relative levels of NDI and ODI in white matter are established early in development and this organizational pattern is retained throughout the lifespan. These observations are particularly interesting in newborns (Kunz et al., 2014), as high NDI and low ODI values in commissural and projection fibers suggest that these tracts have already undergone substantial maturation *in-utero*.

In terms of relative NDI growth rates across tracts, we found trends that differ from findings in infancy and late childhood to adolescence. In infants and toddlers (0-3 year olds), faster maturation is reported in the corpus callosum and corticospinal tracts (Batalle et al., 2017; Dean et al., 2017; Jelescu et al., 2015), while in late childhood to adolescence (8-13 years old), maturation is more rapid in limbic tracts and the superior longitudinal fasciculi (Mah et al., 2017). In early childhood, we find that maturation in the corpus callosum is slower than other bundles, while maturation rates in the corticospinal tracts and superior longitudinal fasciculi are only slightly faster than the rest of the brain. These findings suggest that early childhood might be a transitional period for neurite density development, wherein commissural and projection fibers (i.e. the corpus callosum and corticospinal tracts) are approaching maturity and changes begin to slow, while changes in association fibers such as the superior longitudinal fasciculi commence more rapid increases relative to other tracts in this age range. Our findings of slow maturation in limbic tracts (i.e. the fornix and cingulum bundles) and slow-to-average maturation in the genu and uncinate fasciculi are consistent with the notion that, relative to changes during early childhood, neurite density of limbic tracts increases more rapidly in late childhood (Mah et al., 2017), while increases in NDI in frontal tracts occur more slowly and persist into adulthood (Chang et al., 2015).

In addition to tract-wise maturational trends, we investigated hemispheric differences in cross-sectional and longitudinal tract values. We found small but significant rightward asymmetry cross-sectionally across all white matter tracts for ODI, and all tracts except for the cingulum bundle for NDI, suggestive of greater right hemisphere neurite density and orientation dispersion. Our universal finding of rightward asymmetry of NODDI metrics is interesting, as evidence of white matter asymmetry across the lifespan has been mixed (Bonekamp et al., 2007; Carper et al., 2016; Dean et al., 2017; Takao et al., 2011; Thiebaut de Schotten et al., 2011). Only one study, in newborns, has reported hemispheric asymmetries in NODDI metrics (Dean et al., 2017), finding leftward NDI and ODI asymmetry in the stria terminalis and internal capsule, and rightward asymmetry in the superior corona radiata, superior longitudinal and uncinate fasciculi. These findings differ from those presented here, suggesting that age-related changes in NDI and ODI from birth to early childhood may have a rightward asymmetry to arrive at ubiquitously higher right hemisphere values by approximately age 4, though this hypothesis requires testing. Although there is some conflicting evidence (Krogsrud et al., 2016), most diffusion imaging studies suggest there are few hemispheric differences in white matter maturation from early childhood to adulthood (Genc et al., 2019; Kodiweera et al., 2016; Lebel and Beaulieu, 2011, 2009; Takao et al., 2011), which is consistent with our longitudinal findings. Instead, it appears that hemispheric asymmetries in white matter properties emerge primarily during prenatal and early postnatal periods (Dean et al., 2017; Song et al., 2015). It will be interesting for future studies to investigate hemispheric asymmetries in NDI and ODI, particularly during infancy and toddlerhood, and to determine if these properties are more consistently asymmetric in comparison to other structural features such as those inferred from DTI metrics.

### 4.2 Interpretation in relation to the DTI literature

Developmental increases in NDI have been reported along with increases in FA and decreases in MD from childhood to adulthood (Dubois et al., 2014; Lebel et al., 2012; Westlye et al., 2010). DTI properties depend on multiple white matter structural features, including changes in axonal packing and myelination (Dubois et al., 2014; Lebel and Deoni, 2018; Mukherjee and McKinstry, 2006). Several previous studies have shown that FA and MD correlate with NDI at different ages (Billiet et al., 2015; Mah et al., 2017; Ota et al., 2017; Pines et al., 2019), suggesting that to some degree they reflect similar white matter properties. Interestingly however, the strength of correlations between FA/MD and NDI appears to change with age: in late childhood and early adolescence, stronger and more widespread associations with NDI are found for MD as compared to FA (Mah et al., 2017), whereas in adults, stronger associations are found between NDI and FA than between NDI and MD (Billiet et al., 2015; Ota et al., 2017; Pines et al., 2019). This variability in across-metric correlations with age is likely due to FA and MD being influenced by multiple structural properties that change heterochronously with age (Dubois et al., 2014; Lebel and Deoni, 2018; Mukherjee and McKinstry, 2006). Importantly, several studies have demonstrated that NDI correlates more strongly with age than does either FA or MD (Chang et al., 2015; Genc et al., 2017a; Mah et al., 2017; Ota et al., 2017; Pines et al., 2019), suggesting that NDI may be better suited for tracking neurodevelopmental changes, likely owing to its greater structural specificity. Our own findings support this notion: NDI percentage increases reported here (7.1-10.1%) were more substantial than changes in FA and MD (4.1-7.9% and 3.9-5.9%, respectively) found in a previous study from our group in an overlapping sample (Dimond et al., 2019a).

In previous work, changes in tract-wise FA and MD throughout the lifespan have suggested that maturation occurs earliest in commissural and projection fibers, followed by association fibers, and finally frontal-temporal fibers (Dubois et al., 2014; Hermoye et al., 2006; Lebel et al., 2012; Lebel and Beaulieu, 2011; Westlye et al., 2010; Yoshida et al., 2013). This pattern has recently been shown to hold true for DTI metric changes in early childhood (Dimond et al., 2019a; Reynolds et al., 2019a). In contrast to these DTI findings, as well as NDI changes in infancy (Dean et al., 2015) and late childhood to adulthood (Chang et al., 2015; Mah et al., 2017), we did not observe a clear spatial pattern in tract-wise NDI increases. Taken together, this suggests that neurite density increases follow canonical tract-wise growth trends across the lifespan, though this pattern is not prominent in early childhood. Still, we note some similarities between tract-wise NDI growth reported here, and DTI changes previously reported in an overlapping sample (Dimond et al., 2019a): both NDI and FA increase relatively faster in the arcuate fasciculi, and relatively slower in the cingulum and fornix bundles, while maturation in the superior longitudinal fasciculi is slightly above average in terms of both NDI and MD. Collectively, the similarities and differences in NDI and FA/MD maturational changes throughout the lifespan suggest there is a relative ordering in the maturation of white matter tracts, though individual structural properties follow somewhat unique developmental trajectories.

### 4.3 Biophysical modeling and histological insight into neurite density changes with age

Since its introduction, NODDI (Zhang et al., 2012) has become a popular tool for assessing white matter microstructure, though other biophysical diffusion models have been developed in recent years to estimate neurite or axonal density in-vivo. White matter tract integrity (WMTI; Fieremans et al., 2011) is a two-component model that has been utilized to show developmental increases in axonal water fraction (AWF; the fraction of diffusion signal that is attributed to intra-cellular, as opposed to extra-cellular diffusion, which approximates axonal volume or density) during infancy (Jelescu et al., 2015) and late childhood-adolescence (Huber et al., 2019). Similar to our findings with NDI, Huber et al. (2019) found that AWF increases with age in almost all white matter tracts, with a stronger effect in the arcuate fasciculi where we found NDI changes to be most rapid.

Apparent fibre density (AFD) (Raffelt et al., 2012), calculated from fiber orientation distributions estimated by constrained spherical deconvolution techniques (Dhollander and Connelly, 2016; Jeurissen et al., 2014; Tournier et al., 2007, 2004) and evaluated using the Fixel-Based Analysis (FBA) framework (Raffelt et al., 2017), is another measure of neurite/axonal density that has been shown to increase with age (Dimond et al., 2019a; Genc et al., 2018a, 2018b, 2019). In an overlapping sample, we found FD changes (Dimond et al., 2019a) to be less widespread than changes in NDI reported here, with smaller percentage changes across tracts (3.1-7.1% for FD; 7.1-10.1% for NDI), and no significant growth in the frontal-temporal (e.g. the uncinate and inferior longitudinal fasciculi) or limbic (i.e. the cingulum and fornix bundles) fibers. We also found that rates of change for FD were more variable across tracts, with more rapid commissural/project fiber maturation and slower frontal/temporal tract change, in contrast to the relatively homogeneous tract-wise growth rates found here. These findings might seem contradictory at first sight, though it is important to note that while NDI is calculated at the voxel level, FD is calculated at the level of individually oriented fiber bundle populations within a voxel, called “fixels” (Raffelt et al., 2017). It is possible that more uniform growth and decreased variability in NDI rates of change across tracts may be the result of voxel aggregation, where voxels containing multiple discrete fibre bundles are assigned a single neurite density estimate only, resulting in a shared growth trend amongst those tracts with spatial overlap, and more homogenous maturational trends across tracts.

Our findings suggest that neurite density increases with age, though it is important to note that increases in NDI could potentially be driven by changes in other white matter structural features. NODDI is a multi-compartment model that estimates NDI as the intra-neurite volume fraction relative to the volume of the extra-neurite compartment (Zhang et al., 2012). As such, NDI is potentially sensitive to any microstructural processes that influence the volume ratio of these spatial compartments (Jelescu et al., 2015; Jelescu and Budde, 2017), including: cell genesis (Sigaard et al., 2016), myelination (Williamson and Lyons, 2018), and increases in tract volume (Huang et al., 2006; Vasung et al., 2017) and synaptic density (Huttenlocher, 1979) that have been shown through histological studies to occur during brain development. While estimates of intra-axonal signal fractions from NODDI and other multi-compartment diffusion models have been shown to be generally consistent with histological estimates (Jespersen et al., 2010; Sepehrband et al., 2015), the extent to which specific structural features might be driving maturational changes in NDI remains unclear. Application of additional higher-order models for estimating intra-axonal signal fractions and other specific structural properties (Novikov et al., 2018; Raffelt et al., 2017; Wang et al., 2011, and others, reviewed in Alonso-Ortiz et al., 2015, and Jelescu and Budde, 2017) could be helpful to characterize neurite or axonal density changes with age and to parse out the contributions of other structural changes. Further advancements in biophysical modelling and comparisons to physical measures from histological studies will be important to fully characterize developmental trajectories of specific structural features, such as neurite density, over the course of human brain development.

### 4.4 Cognitive, functional, and neurodevelopmental relevance

During early childhood, cognitive maturation is rapid, with developmental improvements in working memory (Burnett Heyes et al., 2012), math (Garon-Carrier et al., 2018), self-regulation (Montroy et al., 2016), reading (Ferretti et al., 2008), and attention (Breckenridge et al., 2013; Mullane et al., 2016), among other traits. Individual differences in axonal/neurite density have been linked to each of these cognitive traits (Chung et al., 2018; Collins et al., 2019; Genc et al., 2019; Huber et al., 2019; Kelly et al., 2016; Merluzzi et al., 2016), suggesting developmental increases in neurite density may contribute to cognitive-behavioral maturation in early childhood. Considering growth patterns in infancy and late childhood (Batalle et al., 2017; Dean et al., 2017; Jelescu et al., 2015), we interpret our NDI results to suggest maturation during early childhood in association fibers is becoming more rapid, commissural and projection fibers are approaching maturity, and maturation in limbic and frontal tracts remains modest. This may be related to rapid development of skills such as reading and math (Garon-Carrier et al., 2018, Ferretti et al., 2008), nearly complete maturation of sensorimotor processes (Aboitiz and Montiel, 2003; Welniarz et al., 2017), and the beginning of more rapid improvement and refinement of emotional regulation (Ahmed et al., 2015) and executive functions (De Luca and Leventer, 2011), respectively. Our finding that NDI increases were most rapid in the left arcuate fasciculus is particularly interesting, as this tract is known to be important for reading and literacy (Gullick and Booth, 2015; Jess E. Reynolds et al., 2019b; Yeatman et al., 2011) and these neurodevelopmental increases coincide with the rapid development of early literacy and reading skills in early childhood (Ferretti et al., 2008).

Characterizing typical neurodevelopment in early childhood is important as it provides a comparative baseline for detecting abnormal brain development during a period of life which has been associated with the emergence of various neurodevelopmental and psychiatric disorders. Decreased axonal/neurite density has been found in school-aged children who were born very-preterm (VPT) (Kelly et al., 2016), as well as in youth with autism spectrum disorder (ASD) (Dimond et al., 2019b), and in adults with first-episode psychosis (Rae et al., 2017). These structural abnormalities have been associated with decreased IQ and visuomotor function in children born VPT (Kelly et al., 2016; Young et al., 2019), and greater symptoms of social impairment in individuals with ASD (Dimond 2019). These findings, alongside NDI-cognitive associations in healthy individuals (Chung et al., 2018; Collins et al., 2019; Huber et al., 2019; Merluzzi et al., 2016), suggest that neurite density plays an important role in cognitive function, likely mediated by influences on network connectivity. Age-related increases in NDI have been shown to be associated with increased structural network properties (Batalle et al., 2017) and greater functional network connectivity (Deligianni et al., 2016), which is believed to enable faster communication across brain regions in support of various cognitive processes (Zimmermann et al., 2018). Future studies investigating the link between longitudinal changes in neurite density, functional network connectivity and cognitive maturation will be of high interest.

### 4.5 Strengths and limitations

Strengths of the current work include a relatively large sample and the use of mixed effects modeling to leverage longitudinal data. We also implemented several strategies to limit and account for the potential influence of in-scanner head motion on our results, including steps taken prior to and during image acquisition, utilization of recent improvements to DWI preprocessing, data screening, and regression of estimated motion parameters in statistical analyses. Our study is not without limitations, however. Firstly, while we include both males and females, fewer male data were available at the 12-month time point; while this could have led to a female-weighted bias in our estimated trajectories, we note that previous studies found no sex differences in neurite/axonal density or orientation dispersion in childhood to adulthood (Genc et al., 2019, 2018b, 2017b, 2017a; Mah et al., 2017). Similarly, sex does not seem to influence white matter hemispheric asymmetry (Lebel and Beaulieu, 2009; Takao et al., 2011). Secondly, our study did not assess non-linear trends in white matter development. Non-linear trends in NDI and ODI maturation have been reported across the lifespan (Chang et al., 2015), though evidence suggests that linear trends approximate development over short age ranges during childhood (Huber et al., 2019; Mah et al., 2017). This is consistent with our observations here; visualization of data suggests linear fits are most appropriate across major tracts and global white matter within the age range of this cohort.

### 4.6 Conclusions

Early childhood is an important period for brain and cognitive development. Here, we sought to investigate developmental changes in tract-wise neurite density and orientation dispersion in children 4-8 years of age, thereby bridging the gap in the NODDI literature and providing important insights into this critical period of development. We found that NDI increased with age in all major white matter tracts as well as in the global white matter mask, while ODI showed no change. NDI maturational rates did not follow any clear tract-wise pattern, though we note some important tract-wise differences in relation to growth patterns prior to and after early childhood. Our results highlight early childhood as a transitional period for white matter maturation in terms of neurite density, wherein commissural/projection fibers are approaching maturity, maturation in long range association fibers is increasing relative to other tracts in this age range, and changes in limbic/frontal fibers are modest, with the expectation that they will become more rapid towards adulthood. We also found small but ubiquitous rightward asymmetry of NDI and ODI values across tracts, though no asymmetries were found for the longitudinal rates of change. This observation suggests that hemispheric asymmetry of neurite density and orientation dispersion values emerge prior to early childhood. These findings provide insight into tract-wise patterns and developmental trajectories of neurite density during early childhood. Finally, this work provides a comparative basis for assessing abnormal white matter development in neurodevelopmental disorders and has potential implications in relation to cognitive maturation.

## ACKNOWLEDGEMENTS

We thank all the families who participated in this study, as well as staff at the Alberta Children’s Hospital Imaging Centre.

