## Supplementary Table 1 for "Maturation and interhemispheric asymmetry in neurite density and orientation dispersion in early childhood"

|  |  | **Between-Volume Motion** | | | **Within-Volume Motion** | | |
| --- | --- | --- | --- | --- | --- | --- | --- |
|  | TDS | Avg. translation  (mm) | Avg. rotation  (degrees) | Avg. motion  (mm) | Avg. translation  (mm) | Avg. rotation  (degrees) | Avg. motion  (mm) |
| **Mean** | 34.13 | 1.32 | 0.68 | 1.04 | 0.13 | 0.20 | 0.30 |
| **SD** | 25.96 | 0.55 | 0.56 | 0.38 | 0.06 | 0.12 | 0.16 |
| **Min** | 0.00 | 0.66 | 0.14 | 0.57 | 0.07 | 0.09 | 0.15 |
| **Max** | 153.00 | 6.19 | 3.96 | 4.40 | 0.74 | 1.13 | 1.72 |
| **Age association** |  |  |  |  |  |  |  |
| **r-coefficient** | -0.13 | -0.04 | -0.09 | -0.04 | -0.06 | -0.13 | -0.11 |
| **p-value** | 0.08 | 0.60 | 0.25 | 0.63 | 0.41 | 0.08 | 0.15 |

**Supplementary Table 1. Motion metrics and associations with age across the study sample.** Motion metrics of diffusion weighted images for all scans included in the study, calculated as the mean motion across the b=1000s/mm^2^ and b=2000s/mm^2^ shells. Average motion was calculated for each scan as the average translation and rotation, after converting rotation from degrees to millimetres about a 50-millimetre radius sphere. Associations with age were assessed using Pearson correlations conducted in the full study sample. P-values for associations with age are uncorrected for multiple comparisons. TDS=Total dropout slices.
