## Supplementary Table 2 for "Maturation and interhemispheric asymmetry in neurite density and orientation dispersion in early childhood"

|  | Rate of Change  (ΔODI/year) | Predicted Value at Age 4  (ODIx10^-1^) | Predicted Value at Age 8  (ODIx10^-1^) | Percent Increase  from Ages 4-8 (%) |
| --- | --- | --- | --- | --- |
| Right Corticospinal Tract | -7.4x10^-4^ | 3.0 | 3.0 | -9.9x10^-3^ |
| Left Corticospinal Tract | -5.5x10^-4^ | 3.0 | 2.9 | -7.5x10^-3^ |
| Right Sup. Longitudinal Fasciculus | 6.8x10^-4^ | 3.2 | 3.2 | 8.5x10^-3^ |
| Left Sup. Longitudinal Fasciculus | 9.9x10^-4^ | 3.2 | 3.2 | 1.3x10^-2^ |
| Right Inf. Frontal-Occipital Fasciculus | 9.5x10^-4^ | 3.1 | 3.2 | 1.2x10^-2^ |
| Left Inf. Frontal-Occipital Fasciculus | 1.5x10^-3^ | 3.1 | 3.1 | 1.9x10^-2^ |
| Right Inf. Longitudinal Fasciculus | 1.2x10^-3^ | 4.0 | 4.1 | 1.2x10^-2^ |
| Left Inf. Longitudinal Fasciculus | 9.6x10^-4^ | 4.0 | 4.0 | 9.7x10^-3^ |
| Right Arcuate Fasciculus | 6.6x10^-5^ | 3.5 | 3.5 | 7.6x10^-4^ |
| Left Arcuate Fasciculus | 5.8x10^-4^ | 3.4 | 3.5 | 6.8x10^-3^ |
| Right Cingulum | 3.6x10^-4^ | 3.5 | 3.5 | 4.2x10^-3^ |
| Left Cingulum | 5.9x10^-4^ | 3.3 | 3.3 | 7.1x10^-3^ |
| Right Fornix | 1.7x10^-6^ | 2.4 | 2.4 | 2.8x10^-5^ |
| Left Fornix | -2.7x10^-4^ | 2.3 | 2.3 | -4.8x10^-3^ |
| Right Uncinate Fasciculus | 2.2x10^-3^ | 3.6 | 3.6 | 2.5x10^-2^ |
| Left Uncinate Fasciculus | 9.3x10^-4^ | 3.5 | 3.5 | 1.2x10^-2^ |
| Corpus Callosum Genu | 2.1x10^-3^ | 3.4 | 3.4 | 2.4x10^-2^ |
| Corpus Callosum Body | 3.4x10^-4^ | 3.2 | 3.2 | 4.2x10^-3^ |
| Corpus Callosum Splenium | -7.2x10^-5^ | 3.5 | 3.5 | -8.2x10^-4^ |
| Global White Matter Mask | 1.4x10^-3^ | 3.8 | 3.9 | 1.4x10^-2^ |

**Supplementary Table 2: Developmental trajectory values of tract-wise ODI.** ODI did not change significantly from ages 4 to 8 in any major white matter tracts.
